# Comparing the conformational diversity of α1A-Adrenoceptor in Micelles and Phospholipid Bilayer Models

**DOI:** 10.64898/2026.08.10.743833

**Authors:** Mohammad Hossein Tanipour, Feng-Jie Wu, Ashish Sethi, Daniel J. Scott, Paul R. Gooley

## Abstract

α_1A_-adrenoceptor (α_1A_-AR) is a class A G-protein coupled receptor (GPCR) that stimulates smooth muscle contraction in response to adrenaline and noradrenaline. GPCRs exist in a dynamic equilibrium between multiple conformational states. Ligand binding induces structural rearrangements via conserved microswitches, which are thought to shift the equilibrium and trigger signalling. For structural and biochemical studies, GPCRs must be solubilised from the membrane, typically using detergent micelles. However, detergents can disrupt native dynamics of membrane proteins, potentially confounding experimental results. To address this, phospholipid bilayer mimetics such as nanodiscs and saposin nanoparticles (SNPs) have been developed to provide a more native-like environment. Thermostabilised α_1A_-AR serves as a GPCR prototype and can be expressed and isotopically labelled for NMR purposes. To investigate how membrane mimetics influence the conformational diversity of α_1A_-AR, we compared ^1^H ^13^C_3_-HMQC NMR experiments of ^13^CH_3_-Met labelled α_1A_-AR incorporated into either DDM, LMNG, or SNPs, in presence of ligands with varying efficacies. Several methionine residues are positioned near key microswitches, including M203^5.57^, located closed to the G protein binding site. Its resonance has been proposed as a readout of receptor conformational state, shifting with ligand efficacy. Spectra of ^13^CH_3_-Met labelled α_1A_-AR in LMNG closely resembled those in DDM with some temperature-dependent dynamic variation. In contrast, incorporation into SNPs led to a complete loss of M203^5.57^ signal, consistent with an intermediate exchange rate. These findings demonstrate that the membrane environment can profoundly influence conformational dynamics in GPCR NMR studies. Our results highlight the need to carefully consider membrane environment when interpreting NMR data and underscore the value of benchmarking against biologically relevant controls.

## Introduction

G-protein coupled receptors (GPCRs) are the largest group of integral membrane proteins, which essentially participate in all major physiological processes in mammals. Currently, more than 30% of drugs target this superfamily (1-5). It has been hypothesised that GPCRs are in dynamic equilibrium between multiples states such as inactive, pre-active, active, and transducer-bound active separated by energy barriers (6) which allows many GPCRs to have a basal activity even in the absence of endogenous agonists (7). GPCRs must be purified in detergent micelles to enable structure determination (8, 9). Although, these artificial environments can impact the natural dynamics of the receptors a wide range of downstream experiments have been used to characterise structure and function such as NMR (10-14), X-ray crystallography (15, 16), and cryo-electron microscopy (17, 18). Nanometre-scale discoidal phospholipid bilayer mimetic models such as nanodiscs (19-22), saposin nanoparticles (SNPs) (23), and peptidiscs (24, 25) have been introduced to obtain more reliable structural and dynamic characterisation by providing more native-like environments which also improve sample stability and longevity. NMR spectroscopy can be utilised to study the relatively weak interactions of GPCRs with a wide range of compounds (e.g., drugs and partner proteins) for monitoring protein dynamics on a broad timescale from sub-nanoseconds to milliseconds and longer (26-29). Still, the majority of NMR studies on GPCRs have been conducted in detergent micelles (30, 31), while nanodiscs (32-42) and saposin nanoparticles (43) have shown promising data as a superior alternative for micelles. Noteworthy, little is known about how different mimetic models can impact dynamic and conformational diversity of a given GPCR. Direct comparison of the same GPCR in different mimetic systems remains rare, largely due to the technical challenges of producing stable, isotopically labelled receptor preparations in multiple environments. In this context, the thermostabilised α_1A_-adrenoceptors (α_1A_-AR) provides a valuable model (44), enabling isotopic labelling and high-quality NMR measurements (45). This receptor offers a unique opportunity to directly compare the impact of detergents and phospholipid bilayer mimetics on GPCR conformational dynamics.

α_1_-ARs belong to class A GPCRs that comprise α_1A_, α_1B_, and α_1D_ subtypes, which stimulate smooth muscle contraction in response to binding adrenaline and noradrenaline (46-48). Like other GPCRs, α_1A_-AR exists in multiple conformational states in a dynamic equilibrium (6, 7). The ligand binding modulates this equilibrium according to the efficacy of the ligand, with agonists promoting a shift towards active states, while inverse agonists promoting a shift toward inactive states (45, 49-55). The molecular details of GPCR conformational equilibrium are thought to be involved with structural changes to a series of conserved microswitches, including P^5.50^I^3.40^F^6.44^, C^6.47^W^6.48^xP^6.50^, N^7.49^P^7.50^xxY^7.53^, and D^3.49^R^3.50^Y^3.51^ (Ballesteros–Weinstein numbering was used here) (1, 49, 56-59). The microswitches in α_1A_-AR are located near M115^3.41^ and M203^5.57^ (supplementary Figure 1). In a previous NMR study of α_1A_-AR, the impact of ligand efficacy on receptor conformational state, monitored by chemical shift and line-width dependence of the ^13^C^ε^H_3_ resonances of these methionines, was investigated in DDM micelles (45) To obtain a comparison of the conformational diversity of the receptor upon binding to ligands of different efficacies, here we compare 2D ^1^H ^13^C-HMQC experiments of ^13^C^ε^H_3_-Met labelled α_1A_-AR incorporated in different mimetic systems, including LMNG micelles and SNPs. Consistently, a similar chemical shift pattern was observed in LMNG micelles compared to the previous study in DDM (45). However, a slower conformational exchange rate was found in LMNG by observing two distinct resonances for M203^5.57^ indicative of sampling at least two active conformations. We then optimised the incorporation of α_1A_-AR into SNPs, where we found an intermediate exchange rate that emphasised the different dynamic behaviour of the receptor upon binding to ligands of different efficacies from those incorporated in either DDM or LMNG micelles. These significant differences may allow future characterisation of the dynamics of the different ligand-bound states of these receptors.

## Methods

### Protein constructs

Two thermostabilised constructs of human α_1A_-AR were used: α_1A_-AR-A4 and α_1A_-AR-A4 (L312F) as previously described (45). In brief. The α_1A_-AR-A4 variant contains 15 stabilising mutations over the wild-type receptor, including C14S, T15I in the N-terminal, E171V in ECL2, L36^1.42^I, I65^2.43^V, N67^2.47^Y, L80^2.58^M, F86^2.64^Y, G127^3.53^A, G143^4.42^A, W151^4.50^F, F312^7.39^L, N322^7.49^K, P327^7.54^L, and S329^7.56^Y in the TM regions. The α_1A_-AR-A4 (L312F) contains a back mutation of L312^7.39^F relative to α_1A_-AR-A4 variant. To facilitate protein purification, residues S351-V466 were replaced with a 10x-His tag at the C-terminus of α_1A_-AR. In addition, maltose-binding protein (MBP) and green fluorescent protein (GFP) were fused to the N- and C-termini, respectively, and the entire sequence was sub-cloned into the pDS15, a pQE30-derived vector. HRV 3C protease-cleavage sites were placed between fusion proteins (MBP and GFP) and the α_1A_-AR variants. Methionine assignments of ^13^CH_3_ signals previously established in DDM (45) were used for LMNG micelles and SNPs.

Saposin A was expressed using pIQ vector with a 6xHis tag was fused to the N-terminus for purification via metal affinity chromatography. Following cleavage of the His-tag by tobacco etch virus (TEV) protease, the saposin A sequence was SMGSLPCDICKDVVTAAGDMLKDNATEEEILVYLEKTCDWLPKPNMSASCKEIVDSYLPVILDIIKGEMSRPGEVCSALNLCES.

### Protein expression

α_1A_-AR was expressed using *Escherichia. coli BL21-C43 (D*E3) cells and purified as previously described (45). Briefly, Cells were grown at 37 °C with shaking at 225 rpm, and expression was induced by adding 300 μM Isopropyl β-D-1-thiogalactopyranoside (IPTG) once the optical density at 600 nm (OD_600_) reached 0.6-0.9. For ^13^C^ε^H_3_-Met labelling, a modified minimal media was used (60), supplemented with an amino acid mixture containing lysine, threonine, phenylalanine, leucine, isoleucine, valine, and 50 mg/L of ^13^C^ε^H_3_-Met (Cambridge Stable Isotopes). The mixture was added immediately before IPTG induction to inhibit endogenous methionine biosynthesis, while allowing incorporation of the labelled methionine. Cells were harvested after 16-18 h of shaking at 225 rpm and 20 °C.

Saposin A was expressed in *E. coli Rosetta-Gami (DE3) cells*, which facilitate the formation of disulphide bridges. Protein expression was induced with 1 mM IPTG once the OD_600_ reached 0.9, and cells were harvested after 4 h of shaking at 225 rpm and 37 °C.

### Protein purification

The frozen cells were thawed slowly at room temperature, and ice-cold solubilisation buffer (100 mM HEPES, 300 mM NaCl, 1% DDM, 0.12% CHS, 0.6% CHAPS, 10% glycerol, 1 g/L lysozyme, 50 mg/L DNase I, 0.1 mM PMSF, and a tablet of protease inhibitor cocktail, pH 8) was gradually added. The resuspended cells were incubated for 45 min at 4 °C with gently rocking before sonication (Qsonica sonicator, 30% amplitude, 10 s on/20 s off, for 6-8 min). The lysate was then incubated for 1 h at 4 °C with gentle rocking and clarified by high-speed centrifugation (30000 x g, 4 °C, 45 min). TALON metal affinity resin (Takara) was pre-equilibrated in a glass gravity column with equilibration buffer (25 mM HEPES, 200 mM NaCl, 0.05% DDM, and 10% glycerol, pH 8). The filtered supernatant (0.45 μm syringe filter, Merck Millipore) was incubated with the resin for 2 h at 4 °C while rocking. After discarding the flow-through, the resin was washed with wash buffer (25 mM HEPES, 300 mM NaCl, 0.05% DDM, 10% glycerol, and 10 mM imidazole) to remove non-specific binding. The full-length receptor fused to MBP and GPF was eluted with elution buffer containing high concentration of imidazole (25 mM HEPES, 200 mM NaCl, 0.05% DDM, 10% glycerol, and 300 mM imidazole, pH 8). The eluate was concentrated using 100-kDa MWCO centrifugal filter (Amicon ultra, Merck Millipore) and desalted on a PD-10 desalting column (Sephadex G-25M, GE Healthcare) to remove imidazole. MBP and GFP fusions were cleaved overnight at 4 °C using GST-tagged HRV 3C protease in presence of 100 mM Na_2_SO_4_ and 1 mM TCEP (tris(2-carboxyethyl)phosphine) (Sigma-Aldrich). The cleaved solution was incubated with TALON resin for 2 h at 4 °C, and the flow-through containing cleaved GFP, MPB, and 3C protease was discarded. Next, the resin was washed, and the receptor was eluted with buffer containing high concentration of imidazole. For LMNG samples, DDM was gradually exchanged for LMNG by applying wash buffers with increasing LMNG/DDM ratios, until the final solution contained only LMNG, and the receptor was subsequently eluted in LMNG-containing buffer. The eluate was concentrated using 30-kDa MWCO centrifugal filter (Amicon ultra, Merck Millipore) and subjected to size exclusion chromatography (SEC) on a pre-equilibrated Superdex 200 10/300 column (GE Healthcare) with SEC buffer (25 mM HEPES, 200 mM NaCl, and 0.02% DDM or 0.01% LMNG, pH 8) at a flow rate of 0.5 mL/min. Corresponding fractions were pooled, and the NMR samples were prepared by exchanging into NMR buffer (50 mM sodium phosphate, 50 mM NaCl, and 0.05% DDM or 0.01% LMNG, dissolved in 99.9% ^2^H_2_O, pH 7.4).

For saposin A purification, frozen cells were thawed slowly in solubilisation buffer (100 mM HEPES, 300 mM NaCl, 10% glycerol, 50 mg/L DNase I, 0.1 mM PMSF, and a tablet of protease inhibitor cocktail, pH 8). The resuspended cells were lysed using a high-pressure cell crusher (Avestin Emulsiflex, ATA scientific) for 8-12 cycles. The cell debris was removed by centrifugation (16000 rpm, 25 °C, 20 min), after which the supernatant was heated for 10-15 min at 85 °C in a preheated water bath. Precipitated proteins were removed by centrifugation (16000 rpm, 25 °C, 45 min). The resulting supernatant was incubated with pre-equilibrated His60 Ni superflow resin (Takara) in equilibration buffer (25 mM HEPES, 200 mM NaCl, 5 mM imidazole, pH 8) for 2 h at room temperature with gentle rocking. After discarding the flow-through, the resin was washed with wash buffer (25 mM HEPES, 300 mM NaCl, 20 mM imidazole, pH 8). His-tagged saposin A was eluted using elution buffer (25 mM HEPES, 200 mM NaCl, 400 mM imidazole, pH 8). The eluate was concentrated using 3-kDa MWCO centrifugal filter (Amicon ultra, Merck Millipore), and the His-tag was cleaved overnight at 4 °C with TEV protease. The cleaved mixture was heated at 65 °C for 15-20 min to precipitate impurities and TEV protease, which were subsequently removed by centrifugation (16000 rpm, 4 °C, 45 min). The purity of saposin A in the supernatant was assessed by SDS-PAGE.

### Lipid preparation

Fresh phospholipid stock solution of 1-palmitoyl-2-oleoyl-sn-glycero-3-phosphocholine (POPC) (Avanti polar lipids, Inc) was prepared for each reconstitution mixture. POPC (100 mM) was dissolved in chloroform in dark glass tubes. The chloroform was then removed by gently streaming N_2_ gas over the solution until a thin lipid layer formed on the tube wall. To ensure complete removal of chloroform, the tube was placed into a vacuum desiccator for 6-8 h. The dried POPC film was subsequently dissolved in buffer (25 mM HEPES, 150 mM NaCl, 100 mM sodium cholate, and 1% DDM, pH 7.4) to obtain 25-50 mM lipid stock solution.

### SNPs self-assembly with α_1A_-AR variants

The reconstitution mixture consisted of purified saposin A, DDM-solubilised phospholipid, and detergent-purified α_1A_-AR. To optimise α_1A_-AR incorporation efficiency, various molar ratios of α_1A_:saposin A:POPC were screened at different pH values. The optimal condition was determined as a 1:20:250 molar ratio at pH 7.4. The α_1A_-AR concentration was maintained at 6-12 μM, with saposin A and POPC concentrations adjusted accordingly. The mixture was incubated for 2 h on ice. Detergent removal was performed overnight using Biobeads (0.5 g/mL), which had been pre-rinsed with methanol, water, and reconstitution buffer (25 mM HEPES, 200 mM NaCl, pH 7.4), before being added to the reconstitution mixture. After removing the Biobeads, the mixture was further purified by IMAC and SEC under detergent-free condition. To separate loaded and unloaded discs, the reconstitution mixture was subjected to IMAC to capture His-tagged α_1A_-AR, while cleaved saposin A lacking a His-tag remained in the flow-through. The mixture was incubated with TALON resin for 2 h at room temperature, then the flow-through containing empty SNPs and free saposin A was discarded. The resin was washed with buffer (25 mM HEPES, 300 mM NaCl, and 15 mM imidazole, pH 8), and incorporated SNPs were eluted with elution buffer containing high concentration of imidazole in the elution buffer (25 mM HEPES, 200 mM NaCl, and 300 mM imidazole, pH 8). The eluate was concentrated with 100-kDa MWCO centrifugal filter (Amicon ultra, Merck Millipore) and further purified by SEC on a Superdex 200 10/300 columns (GE Healthcare) with SEC buffer (50 mM sodium phosphate, 150 mM NaCl, pH 7.4). Corresponding fractions were pooled, and the NMR samples were prepared by buffer exchange into NMR buffer (50 mM sodium phosphate, 50 mM NaCl, dissolved in 99.9% ^2^H_2_O, pH 7.4) using 100-kDa MWCO centrifugal filter (Amicon ultra, Merck Millipore).

### Analytical ultracentrifugation (AUC)

The size and homogeneity of the samples were analysed using an XL-I analytical ultracentrifuge (Beckman Coulter, Indianapolis, USA) equipped with an AnTi-60 rotor. Protein samples were loaded into the sample compartment of double-sector centrepieces, with buffer in the reference compartments. Radial absorbance data were acquired at 20 °C at a rotor speed of 40000 rpm and a wavelength of 280 nm, with radial increments of 0.003 cm in continuous scanning mode. The sedimenting boundaries were fitted to a model that describes the sedimentation of a distribution of sedimentation coefficients (c(s)) with no assumption of heterogeneity using the program SEDFIT. Data were fitted using a regularisation parameter of p = 0.95, floating frictional ratios, and 100 sedimentation coefficient increments over the range of 0–100 S.

### Size Exclusion Chromatography Multi-Angle Laser Light Scattering (SEC-MALS)

The size and homogeneity of the samples were analysed using an LC-20AD liquid chromatography unit with a DAWN EOS MALS detector (Wyatt Technology Corporation). Separation was performed using SEC-Zenix 300 column at a flow rate of 0.35 mL/min and 25 °C. Data were collected and processed using ASTRA 7.3.2 software (Wyatt Technology), and the size of the samples was estimated based on the elution of 1 mg/mL bovine serum albumin (BSA, Sigma-Aldrich).

### Negative-staining electron microscopy

Negative staining electron microscopy (NS-EM) of SNPs (20-50 μg/mL) was performed on freshly glow-discharged carbon grids. Grids were stained with 1% uranyl acetate and imaged on a Talos L120C 120-kV electron microscope equipped with CETA 4×4k CMOS camera at 73 K. Data were collected at nominal magnification of x73,000, corresponding to a pixel size of 1.9 Å.

### Thermostability

The thermostability of α_1A_-AR incorporated into different mimetic models was assessed using CPM dye (7-Diethylamino-3-(4’-Maleimidylphenyl)-4-Methylcoumarin). As previously described (61), CPM is weakly fluorescent until it reacts with cysteine thiol groups (61), allowing thermal denaturing to be monitored through an increase in CPM fluorescence intensity. Protein samples were mixed with CPM dye in a fluorescence-black cuvette, and fluorescence was recorded on a Cary Eclipse fluorescence spectrometer (Agilent) with excitation at 387 nm and emission at 463 nm. Measurements were collected over a temperature range of 20-90 °C at a heating rate of 2 °°C/min. The melting temperature (T_m_) was calculated by plotting the fluorescence intensity against temperature using Graphpad Prism 7.

### Binding competency

The binding competency of α_1A_-AR was examined after purification in detergent micelles and following incorporation into SNPs. Saturation binding assays were performed using fluorescent BODIPY-FL-prazosin (QAPB) to measure total binding, and in the presence of both QAPB and Prazosin (inverse agonist) to assess specific competition binding between ligands. For saturation assay, 5-10 nM α_1A_-AR was aliquoted in triplicate into 96-well black plates (Greiner Bio-one). Serial dilutions of QAPB (0-100 nM) were added to each row, and plates were incubated for 2 h at 4 °C. Fluorescence intensity of bound QAPB was measured (Ex: 485 nm, Em: 520 nm) using a POLARstar OMEGA plate reader (BMG Labtech, Ortenburg, Germany). Non-specific binding was determined by repeating the assay in the presence of 10 μM prazosin in all wells. Competition assays were similarly performed in triplicate using 96-well black plates (Greiner, Bio-one) with 100 μL receptor solution containing 10 nM QAPB and increasing concentrations of agonist or antagonist. QAPB exhibits increased fluorescence upon binding to the receptor; thus, higher binding is reflected by higher fluorescence intensity. Measurements were taken after 2 h of incubation using POLARstar OMEGA plate reader.

### 2D ^1^H,^13^C-HMQC NMR experiments

NMR experiments were performed using a 3-mm symmetrical Shigemi microtube (Shigemi Inc, Allison Park, PA) to minimise the required sample volume. Ligands at various concentrations were added to the samples at least 30 min prior to data acquisition. Measurements of α_1A_-AR A4 (L312F) in LMNG and SNPs were conducted at 25 °°C and 35 °C on a 700 MHz Avance IIIHD spectrometer equipped with a triple resonance cryoprobe. 2D ^1^H,^13^C SOFAST-HMQC experiments (62) were used to record the ^13^C^ε^H_3_-Met signals, with excitation by a 2.25 ms PC9 120° ^1^H pulse and refocusing by a 1 ms r-SNOB 180° ^1^H pulse. Spectral widths were set to 12 ppm (^1^H) and 25 ppm (^13^C), with a 0.4 s inter-scan delay. A total of 512 x 100 complex points were recorded with 380 scans per FID, corresponding to ∼10 h of acquisition time. 2D ^1^H,^13^C XL-ALSOFAST-HMQC spectra (63) were acquired primarily for LMNG samples, providing enhanced sensitivity by replacing the SOFAST scheme with the ALSOFAST and employing “delayed decoupling” to minimise relaxation losses. Acquisition time was thereby reduced to 4-6 h for 512 x 64 complex points in 256 scans per FID, with a 0.4 s inter-scan delay. Echo/anti-echo gradient coherence acquisition mode was used in XL-ALSOFAST-HMQC, whereas States-TPPI in SOFAST-HMQC. All Spectra were processed using Topspin 3.2 or NMRPipe (64) where data were apodised using a cosine bell window function and zero-filled once in each dimension before Fourier transformation. Spectra were analysed in NMRFAM-Sparky (65).

## Results

### Characterisation of the incorporated α_1A_-AR into SNPs

The optimum incorporation condition for loading SNPs with α_1A_-AR was determined to be a 1:20:250 molar ratio of α_1A_-AR:saposin A:POPC at pH 7.4 (supplementary Figure 2). SNPs are size adjustable, and as previously reported, multiple species can be generated (43). In this study, two main species of α_1A_-AR-loaded SNPs were detected; however, only the smaller species were utilised in the NMR experiments. The overall incorporation efficiency ranged from 40%-60%, decreasing to 20%-50% when considering only the smaller SNPs. Loaded and empty discs were separated by IMAC (Supplementary Figure 3 a), while different loaded SNPs species were further separated by SEC (Supplementary Figure 3 b and c).

AUC analysis comparing the size distribution of loaded SNPs species with empty particles demonstrated a homogenous profile for both empty and smaller loaded SNPs, whereas the larger discs appeared heterogeneous and may contain multiple particle sizes (supplementary Figure 3 d). Although SEC and SEC-MALS indicated that SNPs were generally larger than micelles, the smaller loaded SNPs showed a homogenous monodispersed profile in both analyses (supplementary Figure 4). Visualisation of the smaller loaded SNPs by NS-EM further confirmed their homogeneity and consistent particle size (∼10-12 nm) (Figure 1 a). Importantly, incorporation into SNPs not only preserved receptor architecture but also significantly enhanced thermostability compared with micelles. CPM assays revealed an increase in T_m_ from 42 °C to 58.6 °C for α_1A_-AR upon reconstitution from DDM into SNPs, an improvement that was highly advantageous for conducting NMR experiments at elevated temperatures (Figure 1 b).

**Figure 1.**
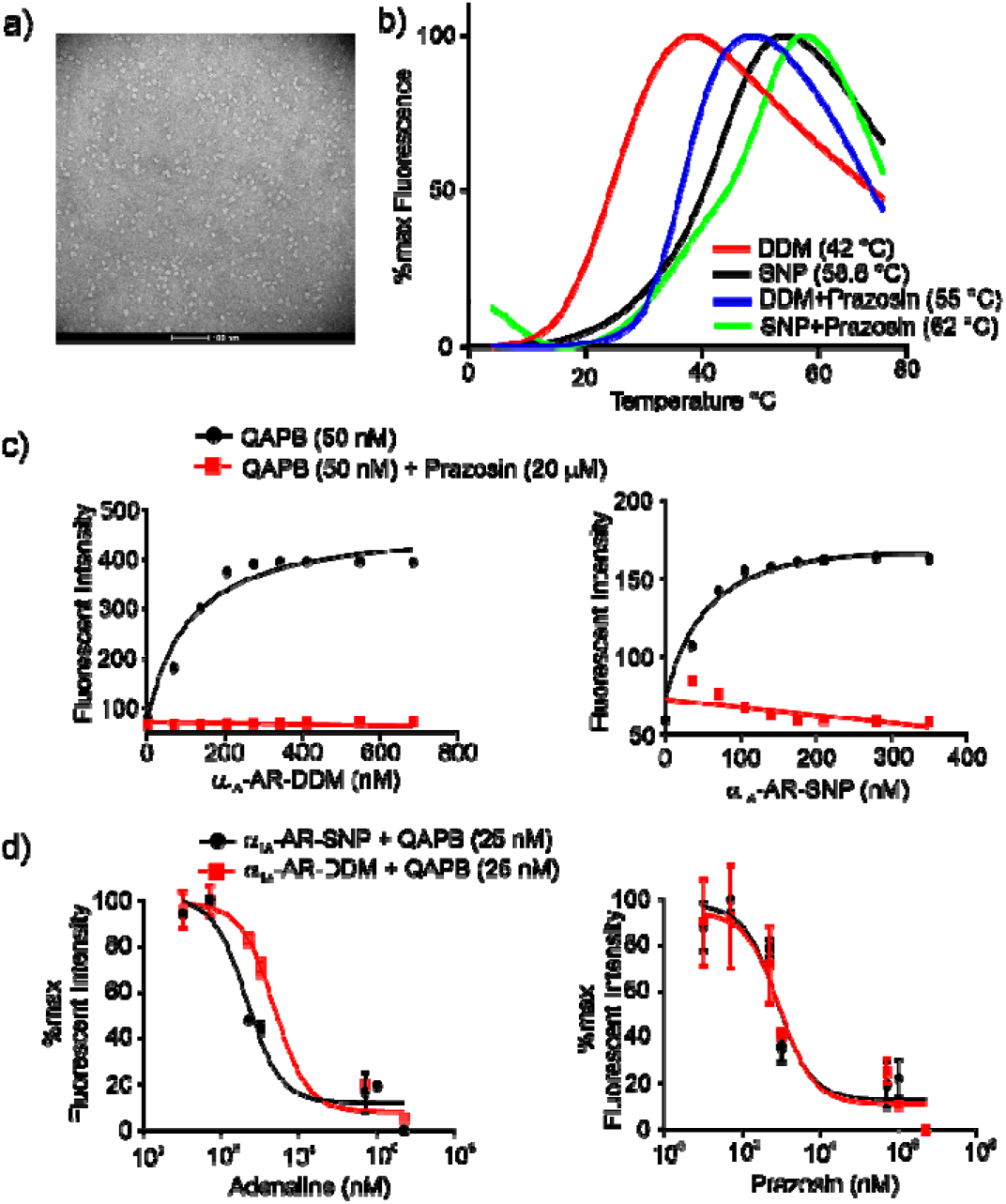
Characterization of α_1A_-AR embedded in SNPs. α_1A_-AR was incorporated optimally into SNP discs at 1:20:250 molar ratio of α_1A_-AR:saposin A:POPC. a) Representative negative-stain micrograph of α_1A_-AR SNPs. b) Comparing CPM thermostability analysis of α_1A_-AR in DDM and SNPs in the apo and inverse agonist (prazosin) bound states. c) Saturation binding assays of α_1A_-AR in DDM and SNPs using QAPB for total binding and QAPB plus prazosin for non-specific binding. d) Competition binding assay of QAPB with adrenaline and prazosin of α_1A_-AR in DDM and SNPs. For DDM sample, the binding buffer contained 0.02% DDM, while no detergent was presented in the SNP assay. All binding data confirm that α_1A_-AR remains well-folded after reconstitution into SNPs. Each data point represents the mean ± SE of triplicate measurements, plotted as fluorescence intensity against α_1A_-AR or competitor concentration. *K*_D_ values were calculated as 110 nM (DDM) and 63 nM (SNPs) using the nonlinear curve fitting algorithm for a single binding site in Graphpad Prism.

Ligand-binding competency assays further confirmed that reconstitution into SNPs did not compromise receptor folding. Loaded SNPs displayed saturation binding patterns for the fluorescent ligand BODIPY-FL-prazosin (QAPB) comparable to those in DDM (Figure 1 c), with calculated *K*_*D*_ of 110 nM (DDM), 63 nM (small SNPs), and 119 nM (large SNPs). Competition binding assay also showed strong agreement between micelles and SNPs: displacement of QAPB by either the agonist adrenaline or the inverse agonist prazosin reduced QAPB fluorescence intensity in a concentration-dependent manner (Figure 1 d).

### Comparing the conformational diversity of α_1A_-AR inserted in DDM or LMNG micelles

α_1A_-AR A4 (L312F) contains six methionine residues: M80^2.58^, M115^3.41^, M145^4.44^, M203^5.57^, M248^ICL3^, and M292^6.55^ (supplementary Figure 1). Of these, M115^3.41^ is located immediately after I114^3.40^ of the transmission switch (PIF motif), M203^5.57^ is positioned above Y125^3.51^ of the DRY motif within the G protein binding pocket, and M292^6.55^ resides in the orthosteric ligand-binding site. Previous NMR studies in DDM revealed that these regions undergo significant ligand-induced local rearrangement between inactive and active conformations (45). In this study, ^13^C^ε^H_3_-Met-α_1A_-AR A4 (L312F) in LMNG was used to record 2D ^1^H, ^13^C-HMQC spectra using XL-ALSOFAST pulse scheme (63) in the presence of ligands of varying efficacies: prazosin (full inverse agonist), WB-4101 (partial inverse agonist), phentolamine (partial inverse agonist), silodosin (neutral antagonist), oxymetazoline (partial agonist), A-61603 (full agonist), and adrenaline (full agonist) (Figure 2 and 3). Methionine assignments were transferred from the previous DDM study (45). Owing to the higher thermostability in LMNG, NMR experiments were performed at both 25 °C (Figure 2) and 35 °C (Figure 3).

**Figure 2.**
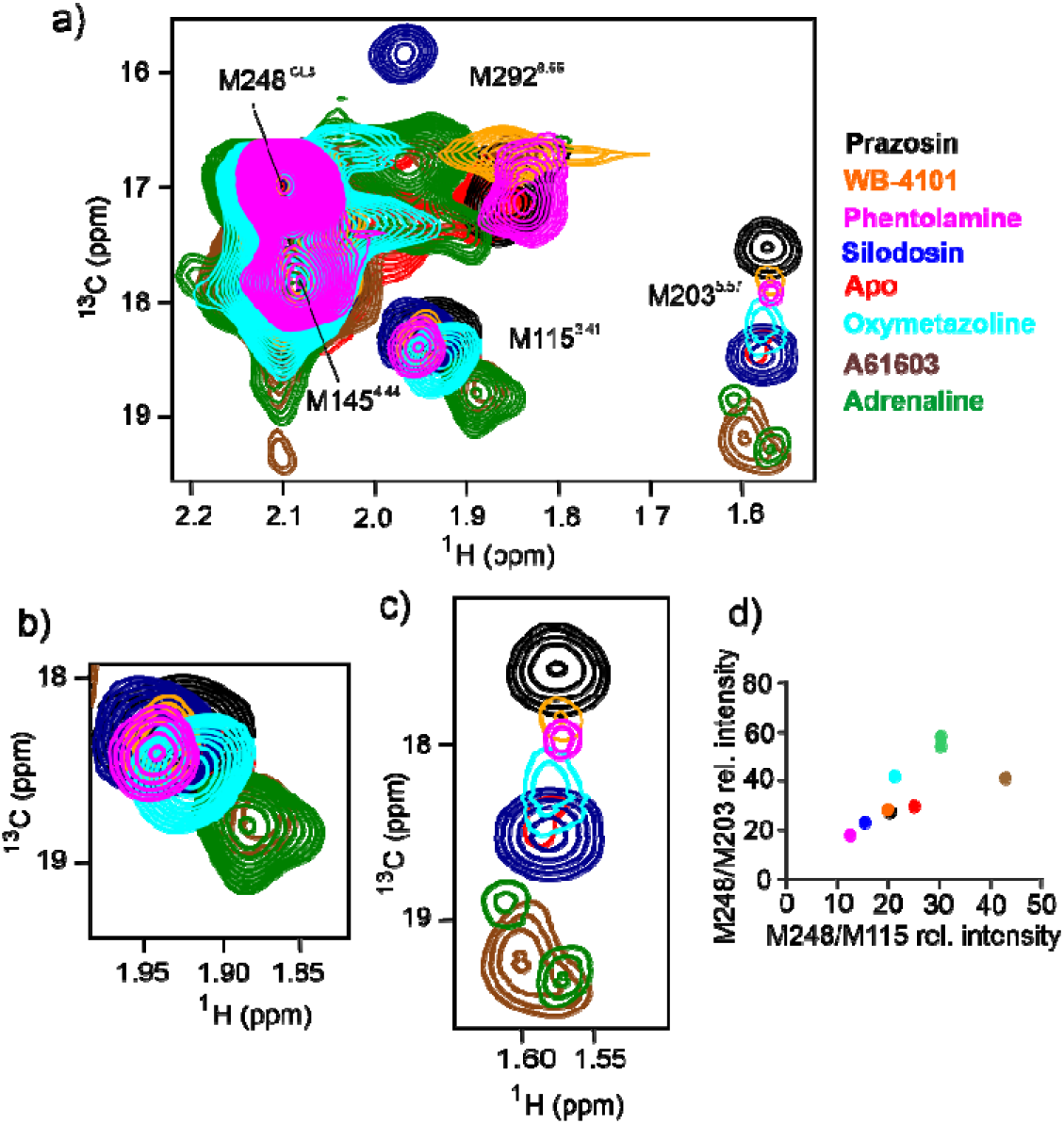
^1^H,^13^C-XL-ALSOFAST-HMQC spectra for ^13^C^ε^H_3_-Met-labelled α_1A_-AR A4 (L312F) in 0.01% LMNG at 25 °C, pH 7.5. a) Overlay of the ^13^C^ε^H_3_-Met region of spectra for the apo state of α_1A_-AR A4 (L312F) (red) and bound to ligands: prazosin (black, inverse agonist), WB-4101 (orange, inverse agonist), phentolamine (magenta, partial inverse agonist), silodosin (blue, neutral antagonist), oxymetazoline (cyan, partial agonist), A-61603 (brown, full agonist), and adrenaline (green, full agonist). b) Expansion of the M115^3.41^ resonance, showing two main resonance clusters corresponding to inactive and active transmission switch states, preferentially occupied by antagonists and agonists, respectively. c) Expansion of M203^5.57^ resonance, showing a linear chemical shift along with ^13^C dimension from inactive to active conformations. Full agonists resonate downfield, while decreasing ligand efficacy toward inverse agonists shift signals upfield. Significant line broadening and splitting of resonances implies slow exchange in LMNG at 25 °C. d) Relative peak intensities of the ^13^C^ε^H_3_ resonances of M203^5.57^ versus that of M115^3.41^, normalised to the reference signal of M248^ICL3^. Agonist binding induced marked signal broadening, while antagonist and inverse agonists reduced conformational heterogeneity, producing sharper peaks than in the apo state.

**Figure 3.**
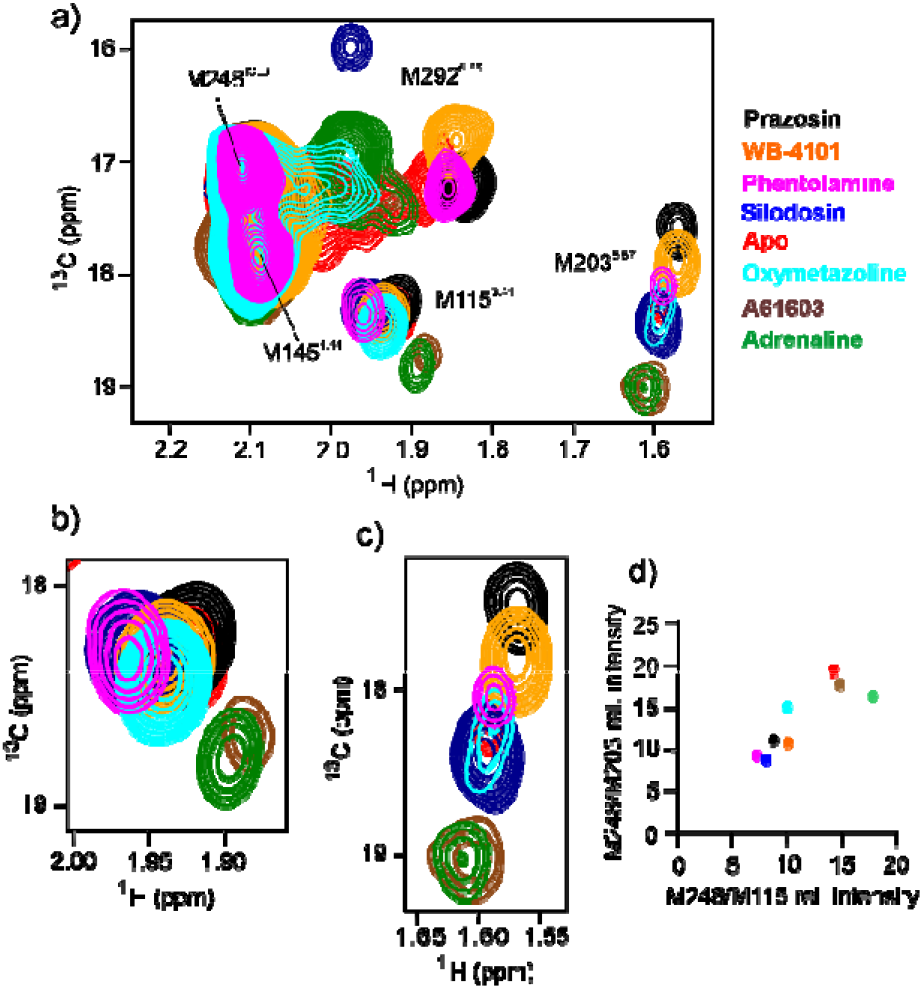
^1^H,^13^C-XL-ALSOFAST-HMQC spectra for ^13^C^ε^H_3_-Met-labelled α_1A_-AR A4 (L312F) in 0.01% LMNG at 35 °C, pH 7.5. a) Overlay of the ^13^C^ε^H_3_-Met region of spectra of α_1A_-AR A4 (L312F) in the apo state (red) and bound to ligands: prazosin (black, inverse agonist), WB-4101 (orange, inverse agonist), phentolamine (magenta, partial inverse agonist), silodosin (blue, neutral antagonist), oxymetazoline (cyan, partial agonist), A-61603 (brown, full agonist), and adrenaline (green, full agonist). b) Expansion of the ^13^C^ε^H_3_ resonance of M115^3.41^, showing two main resonance clusters corresponding to inactive and active transmission switch states, predominantly occupied by antagonists and agonists, respectively. c) Expansion of the ^13^C^ε^H_3_ resonance of M203^5.57^, showing a linear chemical shift along the ^13^C dimension from inactive (upfield) to active (downfield) conformations. A single intense resonance was observed for the M203^5.57^, which implies fast exchange in the LMNG micelles at 35 °C. d) Relative peak intensities of M203^5.57^versus M115^3.41^, normalised to the reference probe M248^ICL3^. At 35 °C, binding of either agonists or antagonists did not induce any signal broadening but instead produced peak sharpening, emphasising an increased exchange rate between conformational states.

At 25 °C, efficacy-dependent ^13^C^ε^H_3_-Met chemical shift changes were observed in LMNG, consistent with the previous DDM study (45). In the apo state, ^13^C^ε^H_3_-Met-α_1A_-AR A4 (L312F) displayed single, well-resolved resonances for each methionine without notable heterogeneity (Figure 2 a), unlike the multiple signals often reported for GPCRs conformational sampling (50, 66). The solvent-exposed M145^4.44^ and M248^ICL3^ served as reference probes (45), displaying intense signals with no ligand-induced changes (Figure 2 a),while M80^2.58^ remained unresolved, likely due to overlap or broadening. M292^6.55^ in the orthosteric pocket exhibited ligand-dependent behaviour similar to DDM. For the different ligands the resonance of M292^6.55^ showed chemical shift differences, reflecting the chemical differences of the ligand. However, under saturating conditions the intensity of the resonance remained unchanged or was slightly more intense. M115^3.41^ displayed two ligand-induced resonance clusters: ∼ 18.3 ppm ^13^C / 1.9 ppm ^1^H for the inverse-agonist (prazosin, WB-4101, phentolamine) and antagonist-bound (silodosin) states and ∼ 18.8 ppm ^13^C / 1.8 ppm ^1^H for agonist-bound states (adrenaline and A61603), with the partial agonist oxymetazoline producing an intermediate shift (Figure 2 a and b). These clusters likely correspond to inactive and active transmission switch conformations. For M203^5.57^, the ^1^H chemical shift was consistently upfield (1.5-1.6 ppm) (Figure 2 a and c) due to the ring-current effect of Y125^3.51^ in the DRY motif (45). As in DDM, (45) the chemical shift of its resonance correlated with ligand efficacy rather than affinity, reflecting equilibria among inactive, intermediate, and active states. Inverse agonists shifted the resonance upfield (inactive states), whereas agonists shifted it downfield (active states), and the neutral antagonist aligned with the apo state (Figure 2 c). The ^13^C chemical shift of the ^13^C^ε^H_3_-M203 group reflects its χ3 rotamers: ∼17.5-18 ppm (±gauche, inactive), ∼18.2 ppm (apo/neutral antagonist), and ∼19-19.4 (trans, active) (45, 67).

Despite the overall similarity in the M203^5.57^pattern, the most prominent difference in the conformational dynamics of the α_1A_-AR A4 (L312F) in LMNG compared with DDM was observed at this residue. In DDM, ^13^C^ε^H_3_ signal of M203^5.57^ exhibited generally a fast exchange regime, appearing as an averaged resonance for inverse-agonist or antagonist-bound states and an intermediate exchange regime for agonist-bound state (45). In LMNG, fast exchange was maintained for the full inverse-agonist, prazosin and the antagonist, silodosin. For partial inverse agonists and partial agonists, the resonance shows intermediate exchange, and for full-agonist, especially adrenaline, the resonance splits into two signals (18.8 and 19.3 ppm for ^13^C) consistent with a slow exchange regime (Figure 2 a and c), suggesting two distinct active states. Raising the temperature to 35 °C significantly affected receptor dynamics where all resonances generally sharpen becoming more intense (Figure 3, Supplementary Table 1 and 2). Most importantly, the resonances of adrenaline collapse to single resonance reflecting a fast exchange regime. While less pronounced, the ^13^C^ε^H_3_ signal of M115^3.41^ also showed a similar chemical shift and broadening dependence on ligand efficacy where agonists appear to increase conformational heterogeneity and promote transitions toward active states. Consistent with previous findings in DDM (45) analysis of the chemical shift difference (Δδ) of the M203^5.57^ (Supplementary Figure 5) revealed a linear correlation with ligand efficacy and no correlation with ligand affinity. These findings suggest that ligand-dependent modulation of the transmission and DRY microswitches is preserved in LMNG, with a temperature-dependent behaviour in exchange kinetics. The observed behaviour further supports maintained allosteric coupling between the orthosteric and G protein binding site in α_1A_-AR A4 (L312F).

### Comparing the conformational diversity of α_1A_-AR embedded in micelles and phospholipid bilayers

In an effort to reproduce a bilayer environment ^13^C^ε^H_3_-Met-α_1A_-AR A4 (L312F) purified in micelles was reconstituted into SNPs. Owing to the significantly higher thermostability of α_1A_-AR in SNPs (Figure 1b), 2D ^1^H,^13^C-SOFAST-HMQC experiments were acquired at 35 °C (Figure 4). Overall, the spectra of α_1A_-AR A4 (L312F) in SNPs suggested altered conformational diversity compared with micellar models. As in micelles, M80^2.58^ was not detected, while the solvent-exposed probes M248^ICL3^ and M145^4.44^ produced intense resonances at chemical shifts similar to those in DDM and LMNG (Figure 4 a). These signals were unaffected by ligand binding, suggesting that solvent exposure was preserved in the phospholipid bilayer environment and validating their use as internal references. The presence of a clear ^13^C^ε^H_3_ signal for M292^6.55^ in the apo state suggested that incorporation into relatively larger SNPs (in comparison to micelles) did not markedly affect linewidth. Its chemical shift behaviour in the orthosteric pocket closely resembled that in micelles, confirming that receptor folding was maintained after SNPs reconstitution. Inverse agonists (prazosin, WB-4101, phentolamine) shifted M292^6.55^ signal upfield in ^1^H dimension relative to the apo state, whereas the partial agonist oxymetazoline and full agonists (A61603, adrenaline) caused pronounced line broadening and downfield shifts (Figure 4 a).

**Figure 4.**
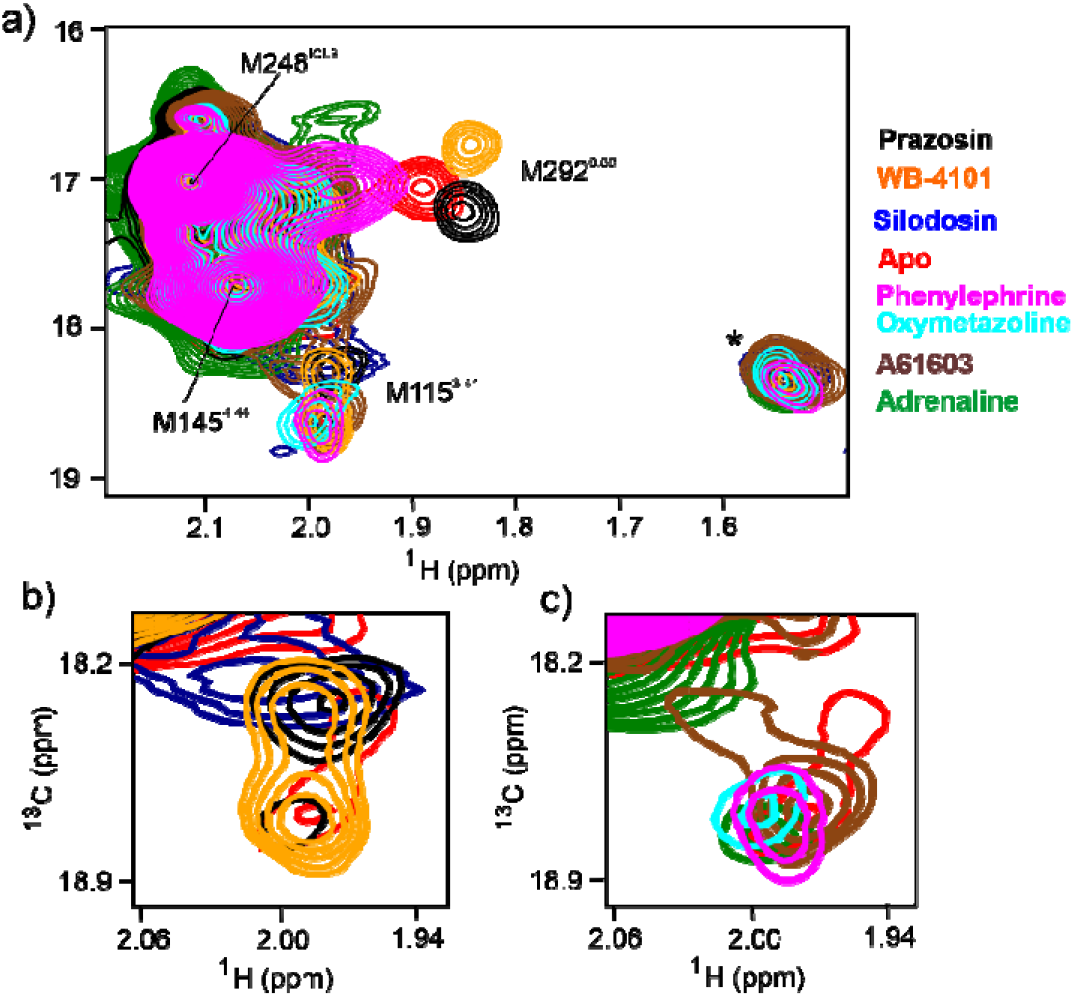
^1^H-^13^C SOFAST-HMQC spectra of ^13^C^ε^H_3_-Met-labelled α_1A_-AR A4 (L312F) incorporated into SNPs at 35 °C, pH 7.5. a) Overlay of the ^13^C^ε^H_3_-Met region of spectra of α_1A_-AR A4 (L312F) in the apo state (red) and bound to ligands: prazosin (black, inverse agonist), WB-4101 (orange, inverse agonist), silodosin (blue, neutral antagonist), phenylephrine (magenta, partial agonist), oxymetazoline (cyan, partial agonist), A-61603 (brown, full agonist), and adrenaline (green, full agonist). A single resonance is observed for M292^6.55^ where its chemical shift is dependent on the nature of the ligand. M115^3.41^ shows efficacy dependent shifts similar to those in LMNG and DDM. Significant line broadening of M203^5.57^ resulted in disappearance of its signal(s), which implies motion on an intermediate timescale at the G protein binding site of the receptor incorporated into SNPs. A non-methionine background signal is marked (*). b) and c) Expansion around the ^13^C^ε^H_3_ resonance of M115^3.41^ from panel a). The apo resonance is split into two weak signals. In b) shifts by inverse agonists and antagonists show a trend for selection of a state corresponding with the upfield signal of the apo state. In c) shifts by agonists show the resonances that align with a state corresponding to the downfield signal of the apo state.

Unlike the solvent-exposed methionines, significant line broadening was observed for the efficacy-sensing probes: M115^3.41^ (near the transmission switch) and M203^5.57^ (close to the G protein binding site). For M115^3.41^, two signals were detected for the ^13^C^ε^H_3_ in the apo state with one signal upfield at 1.8 and 18.3 ppm (^1^H, ^13^C) and the other downfield at 2.0 and 18.7 ppm (^1^H,^13^C) (Figure 4). Incorporation into SNPs produced a slower exchange regime in the transmission switch, as indicated by two distinct resonances, but the chemical shift of ^13^C^ε^H_3_ of M115^3.41^ remained consistent with those in LMNG and DDM. Interestingly, addition of the inverse agonist prazosin shifted the ^13^C^ε^H_3_ resonance of M115^3.41^ to the peak in the upfield position, with loss of the downfield signal, suggesting this upfield resonance corresponds to the most inactive conformation (Figure 4 b). In contrast, agonist binding (adrenaline, A61603) shifted the ^13^C^ε^H_3_ signal to the peak in the downfield, consistent with an active state (Figure 4 c). These data suggest that the two signals of the ^13^C^ε^H_3_ resonance of M115^3.41^ represent an equilibrium between inactive and active (or preactive) transmission switch conformations (6). As expected, partial inverse agonists WB-4101 and neutral antagonist silodosin did not mimic prazosin, instead producing two ^13^C^ε^H_3_ resonances consistent with a slow exchange regime (Figure 4 b). Partial agonist (oxymetazoline, phenylephrine) shifted the ^13^C^ε^H_3_ resonance of M115^3.41^ to intermediate positions between inverse agonist- and agonist-bound states, in line with their partial efficacies (Figure 4 c).

No resolved resonance was detected for the ^13^C^ε^H_3_ signal of M203^5.57^ in α_1A_-AR A4 (L312F) reconstituted into SNPs, either in the apo state or in the presence of agonists or antagonists, most likely due to severe line broadening. This behaviour suggests a conformational exchange on an intermediate timescale regime at the G protein binding site (DRY motif), indicating a slowing the dynamic heterogeneity in the bilayer environment compared with micelles.

## Discussion

Over the past decades, NMR studies of diffusible ligand-activated GPCRs have provided important insights into how ligands modulate the conformational landscape of these receptors (68). These studies suggest that GPCRs exist in dynamic equilibria between inactive, intermediate, and active states, which can be shifted by ligands of varying efficacies (33, 50, 68-70). Intramolecular communication between conserved microswitches is proposed to govern these structural transitions, with ligand-specific modulation reflected in changes to NMR chemical shift and/or signal intensities (67, 71).

Detergent micelles, particularly DDM and LMNG, has been widely used for GPCR characterisation (68, 72, 73). LMNG, with two hydrophobic chains, perhaps more closely mimics the architecture of a lipid than the single chain DDM (74), as demonstrated by its lower monomer off rate (32) and lower CMC (0.001% vs 0.0165%) (75, 76). Molecular dynamics simulations further showed that LMNG enhances aliphatic chain density around the hydrophobic regions of A_2A_AR and forms more hydrogen bonds between head groups (77). LMNG polar groups form persistent bifurcated hydrogen bonds between the TM helices of β_2_-AR, reducing flexibility but increasing stability. Consistent with this, NMR studies of LMNG reconstituted β_2_-AR were conducted at higher temperatures than in DDM, where LMNG’s lower off-rate facilitated detection of distinct functional conformations (32). Despite the practical advantages of micelles, including small size and ease of preparation (11, 78, 79), concerns remain regarding their ability of preserve native protein function. Micelles may perturb folding and disrupt GPCR-G protein interaction, due to poorly mimicking the lipid bilayer (8, 10, 80). To address these limitations, bilayer systems such as nanodiscs (20), peptidiscs (24), and saposin nanoparticles (SNPs) (23) have emerged as superior native-like environments. These systems enhance protein stability, permit experiments at elevated temperatures, and potentially provide more physiologically relevant conditions for dynamic studies. Moreover, their capability to stabilise ternary GPCR-G protein complexes further underscores their value in structural investigations (81).

The previous NMR study of α_1A_-AR in DDM investigated its conformational diversity upon ligand binding (45). To obtain more reliable data on ligand-induced conformational alterations, we employed both LMNG micelles and phospholipid bilayer systems (SNPs). SNPs have been previously utilised for the structural studies of membrane proteins, including cryo-EM (82, 83), FRET (84), Mass Spectrometry (85), and NMR (43). Here, reconstitution of α_1A_-AR into SNPs was optimised at 1:20:250 molar ratio of α_1A_-AR:saposin A:lipid at pH 7.4. Compared with micelles, these SNPs are larger particles, as characterised by NS-EM, SEC, AUC, and SEC-MALS, yet the protein incorporated SNPs remained highly homogenous (Figure 1, Supplementary Figure 2 and 3). The SNPs provided a stable environment for α_1A_-AR, reflected by increased longevity and thermostability (Figure 1). The larger size of SNPs relative to micelles, however, is expected to lead to faster decay of the NMR signal (86), therefore in most phospholipid bilayer models, amino acid deuteration is typically required to obtain high-quality 2D spectra (33). Nevertheless, despite their size, and unlike several GPCR NMR studies in nanodiscs that relied on deuteration (32-34, 36-39), we were able to achieve reasonable ^1^H,^13^C-HMQC spectra quality for ^13^C^ε^H_3_-Met labelled α_1A_-AR without deuteration. This observation is consistent with previous SNP studies reported for non-deuterated ^1^H,^15^N OmpX and ^13^C^ε^H_3_-Met β_1_-AR (43).

In the DDM study of α_1A_-AR (45), three methionine probes were highlighted: M115^3.41^, reporting conformational alteration of the transmission switch (PIF motif); M203^5.57^, probing the DRY motif or G protein binding site; and M292^6.55^, positioned in the orthosteric ligand-binding pocket. M292^6.55^ directly senses interactions with ligand chemical structures, whereas M115^3.41^ and M203^5.57^ indirectly report on conformational changes within the microswitches (Supplementary Figure 1). Our NMR experiments in LMNG (Figures 2 and 3) reproduced the chemical shifts of all methionine probes, including the conformationally sensitive probes, M115^3.41^, M203^5.57^, and M292^6.55^ consistent with data previously reported in DDM at 25 °C.

At 25 °C, the ^13^C^ε^H_3_ resonances of M115^3.41^ and M203^5.57^ varied in chemical shift and linewidth in response to ligand efficacy but not affinity (Figure 2) (45). In DDM (45), M203^5.57^ appeared as a single distinct and relatively intense peak in both apo and inverse agonist or antagonist-bound states, likely reflecting an averaged population exchanging on a fast timescale. In the presence of agonist the resonance broadened, suggesting intermediate exchange between multiple states. Indeed in LMNG, in the adrenaline-bound state (Figure 2c) two slow exchanging distinct peaks were detected for M203^5.57^, providing further evidence of multiple active (or preactive) conformations. A similar phenomenon has been reported for M_2_R labelled with ^13^C^ε^H_3_-Met in LMNG/CHS at 25 °C (87), where agonists induced two distinct chemical shifts for this resonance of M202^5.54^ (homologous to M203^5.57^ in α_1A_-AR) which was interpreted as the co-existence of two conformational states in slow exchange, reflecting distinct agonist-stabilised structural environments. The occurrence of two ^13^C^ε^H_3_ resonances in antagonist-bound states has also been reported for several GPCRs, including β_2_-AR (33, 50, 88), β_1_-AR (89) across micelles and bilayer mimetics (DDM, LMNG, nanodiscs). In these studies, β_1_-and β_2_-AR were found to exist in two slow-exchanging inactive states, with the ^13^C^ε^H_3_ signal of M82^2.53^ in β_2_-AR and M90^2.53^ in β_1_-AR displaying two resonances in the apo and/or antagonist-bound states. These probes lie near the “toggle switch” residue W^6.48^ of the CWxP and PIF motif within the transmission region. Because of their distance (7-9 Å) from the orthosteric pocket, their chemical shifts are more likely to reflect conformational changes in the transmission region rather than direct ligand interaction. Furthermore, a ^19^F-NMR study of BTFMA-labelled A_2A_AR at position V229^6.31^C (90) revealed an ensemble of two inactive and two active states in equilibrium. The inactive states were enriched by inverse agonists and exchanged on the millisecond timescale, consistent with prior β_2_-AR and β_1_-AR NMR studies. The two active states were stabilised by partial or full agonists, supporting a conformational selection mechanism whereby partial agonism arises from preferential sampling of a “less open” active state with G protein coupling capacity.

The chemical shift positions of the ^13^C^ε^H_3_ of M203^5.57^ in α_1A_-AR (L213F) suggests efficacy-dependent equilibrium shifts, from an average of χ3 ±gauche/trans conformer in inverse agonist-bound or inactive states to trans conformers in agonist-bound or active states. Similar ligand efficacy dependence at this position has been reported in β_1_-AR and M_2_R (87, 91). In ^15^N labelled thermostabilised β_1_-AR, the ^1^H,^15^N chemical shifts of Val226^5.57^ (homologous to M203^5.57^ in α_1_-AR) fell along a “straight line” correlating with ligand efficacy. The data collected suggest that the observed conformational changes in the ^13^C^ε^H_3_ resonances of M203^5.57^ reflect the bending of TM5 towards the active conformation. The appearance of multiple states for M203^5.57^ in the adrenaline-bound state of α_1A_-AR(L213F) in LMNG at 25 °C differs from other GPCR studies, such as β_2_-AR, where multiple inactive states were observed for M82^2.53^(33, 50). This discrepancy may reflect the thermostabilised nature of α_1A_-AR A4 (L312F), which is signalling-incompetent due to the N322^7.49^K mutation in the NPxxY motif. This stabilising mutation is proposed to form a salt bridge with D72^2.50^ (45), favouring the inactive state and resulting in a single intense resonance for M203^5.57^ in apo or antagonist-bound states. Raising the temperature from 25 to 35 °C in the ^1^H,^13^C-HMQC experiments remove the line broadening and signal splitting that is observed for the ^13^C^ε^H_3_ signals of M115^3.41^ and M203^5.57^ in agonist-bound states, resulting in single resonances. The overall chemical shift patterns of all probes, however remained the same (Figure 3).

The 2D ^1^H,^13^C-HMQC spectra of α_1A_-AR A4 (L312F) in SNPs suggested altered conformational diversity compared with LMNG and DDM. The similar behaviour of M292^6.55^ in SNPs and micelles, together with binding competency data, confirmed that receptor structure was preserved in SNPs and linewidth was not significantly impacted on this resonance by the larger size of the particle. However, two main differences were found in the ^13^C^ε^H_3_-Met probes of α_1A_-AR A4 (L312F) in SNPs compared with micelles. First, no signal was observed for M203^5.57^ (Figure 4), suggesting that the region around the DRY motif experiences motion on an intermediate timescale in α_1A_-AR. This is consistent with ^19^F-NMR study of β_2_-AR (35) , which showed higher basal activity in lipid bilayer environments than in micelles, suggesting dynamic DRY motif conformations that may underlie the behaviour of M203^5.57^ in α_1A_-AR. Second, M115^3.41^ appeared as two distinct broadened resonances in the apo state in SNPs, rather than the single intense peak observed in LMNG and DDM, indicating slowing of motion near the transmission switch (Figure 4). The two peaks of M115^3.41^ imply that the incorporated receptor into SNPs co-exists in at least two slow-exchanging conformations. This behaviour parallels observations of M112^3.41^ in M_2_R (homologous to M115^3.41^ in α_1A_-AR) in LMNG (87) and M82^2.53^ in β_2_-AR (50), which reflected sampling of multiple PIF motif conformations (PIF_off1_, PIF_off2_,PIF_on_). In α_1A_-AR, the two M115^3.41^ resonances may represent PIF_off1/2_ and PIF_on_. This hypothesis was further supported when agonists shifted the signal to the downfield ^13^C^ε^H_3_ resonance (Figure 4 c), suggesting PIF_on_, while inverse agonists shifted it to the upfield ^13^C^ε^H_3_ signal (Figure 4 b), consistent with PIF_off1/2_ mode. Partial agonists and partial inverse agonists produced intermediate positions, indicating mixed populations of inactive and active states. Together, these data suggest that M115^3.41^ in SNPs samples multiple conformations: inactive (inverse agonist), intermediates/ pre-active (partial agonists/partial inverse agonist), and active (full agonist).

As discussed, the α1A-AR A4 (L312F) variant is signalling-incompetent. Signalling competency was previously restored in the α_1A_-AR A4 (active) variant by introducing back mutations, including K322^7.49^N to recover the NPxxY motif (45). In this active variant in DDM, two ^13^C^ε^H_3_ resonances of M115^3.41^ were detected, consistent with our observation in α_1A_-AR A4 (L312F) in SNPs. These two resonances suggested slow exchange (> ms) between two states, attributed to the restored NPxxY motif. Our data support the interpretation of that study, where the upfield resonance of M115^3.41^ in apo α_1A_-AR-A4(active) was proposed to represent a fully inactive state; this was reinforced by our finding that inverse agonist binding shifted M115^3.41^ predominantlyto the upfield resonance. Furthermore, in the α_1A_-AR A4 (active), M203^5.57^ exhibited marked broadening especially in the agonist-bound states (45), consistent with our observations for M203^5.57^ of α_1A_-AR A4 (L312F) in SNPs. It was assumed that restoration of the NPxxY motif enabled the DRY motif to more willingly sample active states. Taken together, it can be hypothesised that the incorporation of α_1A_-AR A4 (L312F) into SNPs impacts the dynamics of the receptor in a manner resembling the signalling-competent α_1A_-AR A4 (active). This hypothesis agrees with reports of enhanced basal activity of β_2_-AR in phospholipid bilayers compared with micelles (35).

To conclude, our NMR study highlights marked differences in the conformational dynamics of α_1A_-AR across mimetic models, centred on the conserved PIF and DRY microswitches. While nanodisc-based studies typically require deuteration due to poor spectral quality in large bilayer systems, we obtained reasonable 2D NMR spectra in SNPs without deuteration, although the highest spectral resolution was achieved in LMNG. Consistent with other GPCR studies, however, we observed ligand efficacy-dependent modulation of α_1A_-AR conformational equilibria, with slower exchange rate in SNPs. A more complete understanding of these dynamic transitions will require reconstitution of signalling-competent α_1A_-AR variants into SNPs in complex with G proteins to capture the full pathway from inactive to fully active states.

