## Supplementarry for "Comparing the conformational diversity of α1A-Adrenoceptor in Micelles and Phospholipid Bilayer Models": Supplementary_Figures_Comparing the conformational diversity of α1A-Adrenoceptor in Micelles and Phospholipid Bilayer Models.docx

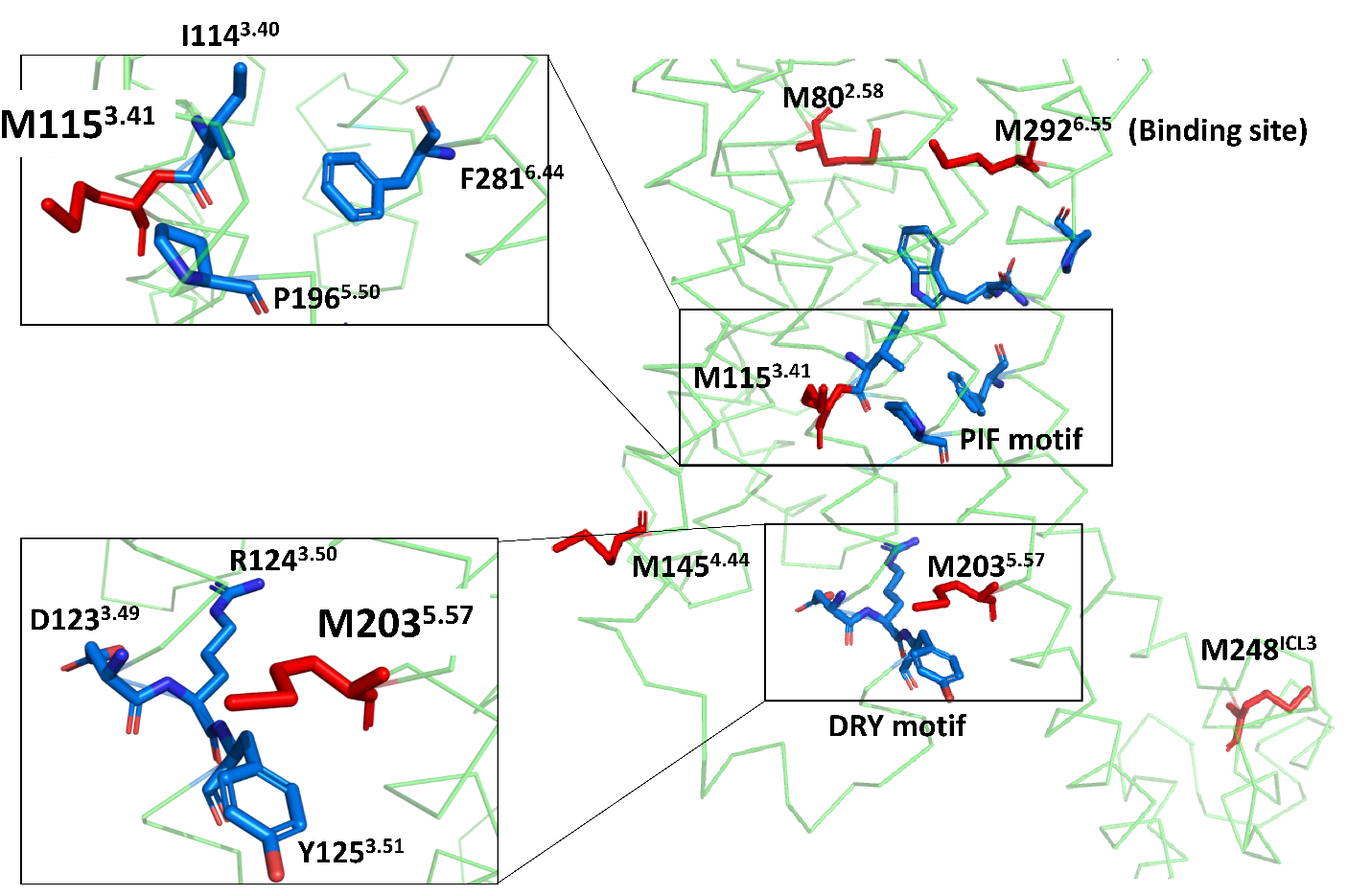

**Supplementary Figure 1**. **Positioning of methionine residues (red) relative to conserved microswitches (blue) in α_1A_-AR A4 (L312F) construct.** α_1A_-AR A4 (L312F) contains six methionine residues: M145^4.44^ and M248^ICL3^ are solvent-exposed; M292^6.55^ resides in the orthosteric binding pocket; M115^3.41^ is adjacent to the I114^3.40^ of the conserved PIF motif; and M203^5.57^ is positioned above the aromatic ring of Y125^3.51^ of the DRY motif.

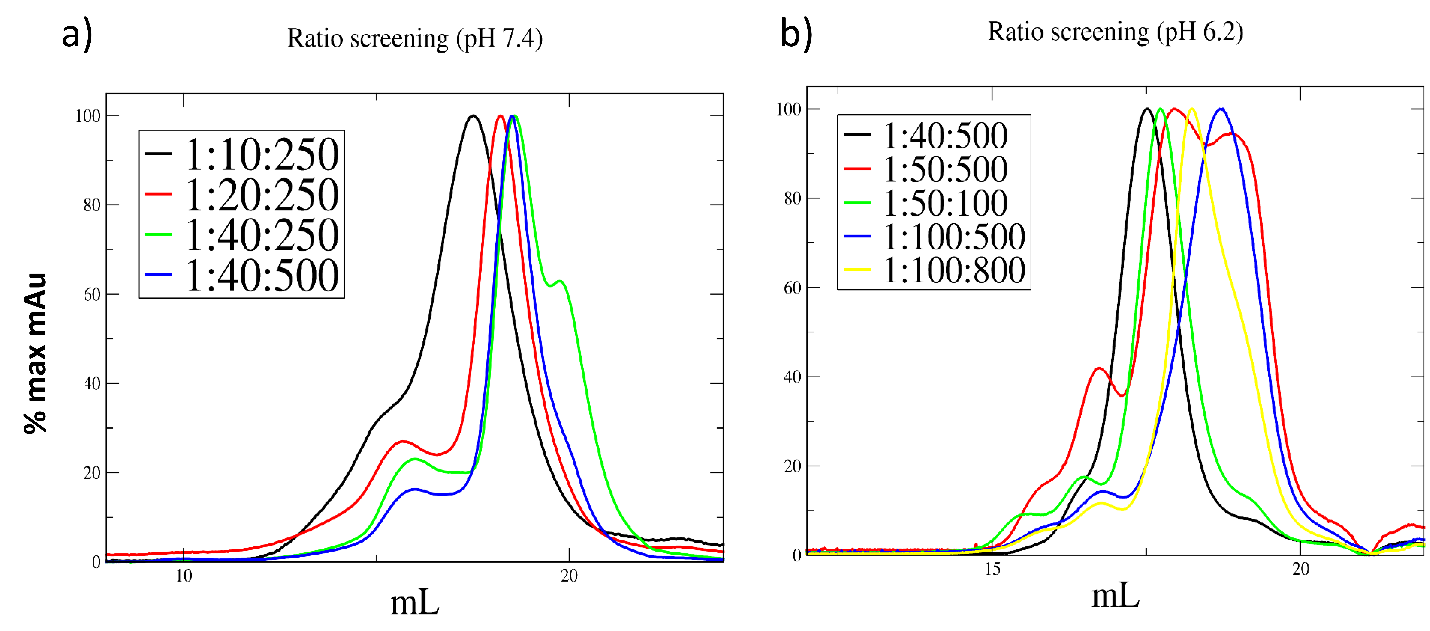

**Supplementary Figure 2**. **Comparing SEC chromatograms of different α_1A_-AR:saposin A:POPC molar ratios.** SEC of α_1A_-AR in SNPs at (a) neutral (pH 7.4) and (b) acidic (pH 6.2) conditions to optimise SNPs assembly. Various reconstitution ratios were examined and analysed using a Superose 6 10/300 SEC column. Higher saposin A ratios (e.g., 1:40:250 (α_1A_-AR:saposin A:POPC) at pH 7.4, and 1:100:500 at pH 6.2) yielded longer retention time, consistent with the formation of smaller, predominantly empty discs. By contrast, lower molar ratios (e.g.,1:10:250, and 1:20:250 at pH 7.4 and 1:40:500 at pH 6.2) produced larger single peaks indicative of greater proportion of loaded discs among assembled nanoparticles.

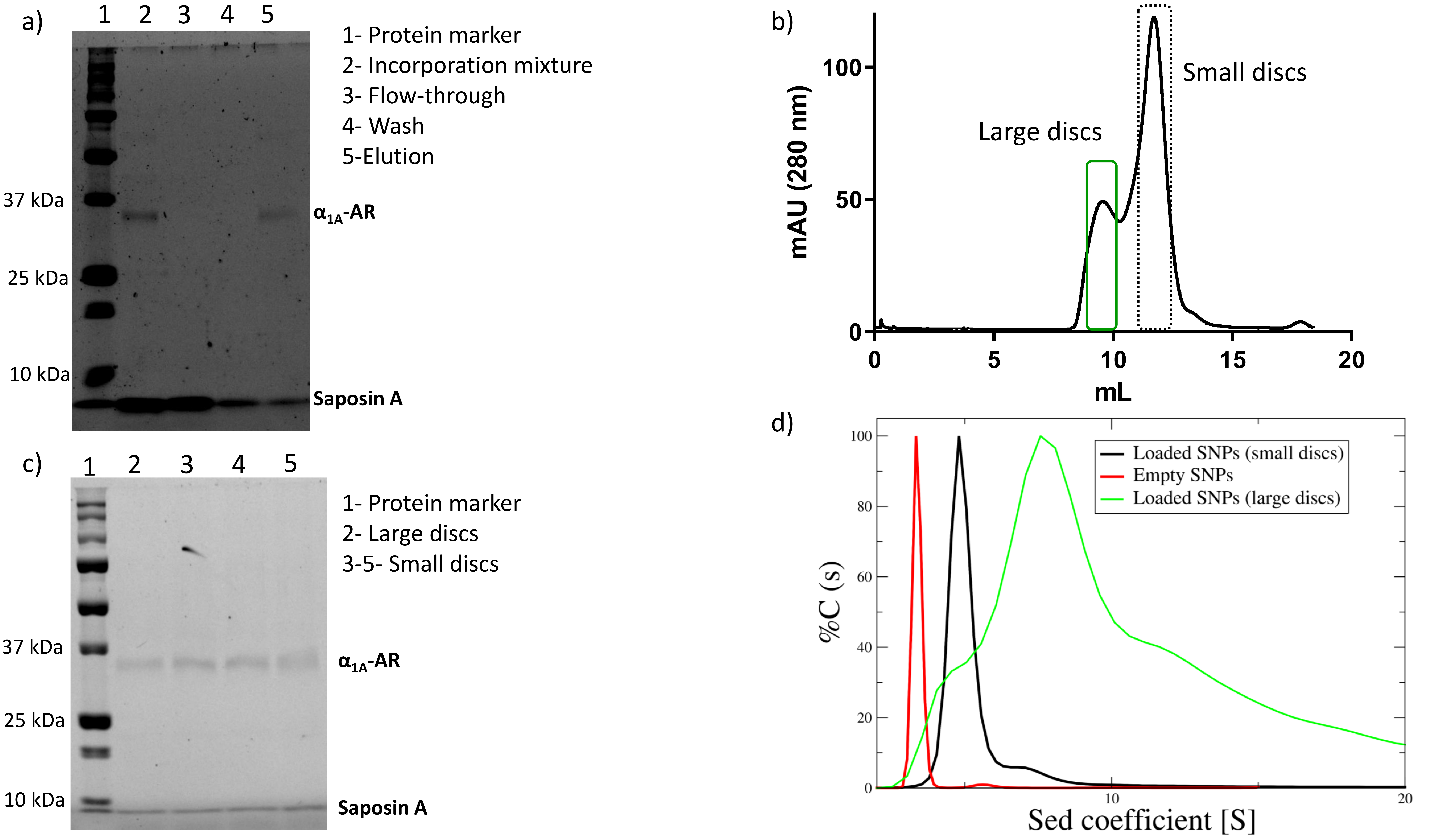

**Supplementary Figure 3**. **Characterization of α_1A_-AR embedded in SNPs.** a) SDS-PAGE for IMAC separation of empty and loaded SNPs. The incorporation mixture contained both α_1A_-AR and saposin A bands (lane 2). Only saposin A was detected in the flowthrough (lane 3), indicating successful separation of empty SNPs. Non-specifically bound empty discs were removed by washing with 20 mM imidazole (lane 4). Elution contained both α_1A_-AR and saposin A bands, confirming purifying of loaded SNPs (lane 5). b) SEC profile (on a Superdex 200 column) of IMAC-eluted SNPs (produced from 1:20:250 molar ratio of α_1A_-AR:saposin A:POPC at pH 7.4) showing two peaks, consistent with distinct species of incorporated discs. The main peak (dotted box) corresponds to smaller loaded SNPs, while a minor peak (green box) represents larger discs. c) SDS-PAGE of SEC fractions: lane 2, large discs (first SEC peak); lanes 3-5, small discs (main SEC peaks. Both show bands for saposin A and α_1A_-AR. d) AUC characterisation of empty (red), small (black), and large(green) loaded SNPs taken from (b). Empty and small loaded SNPs displayed homogeneous distributions, while large loaded SNPs showed heterogeneity with a broad peak and multiple shoulders.

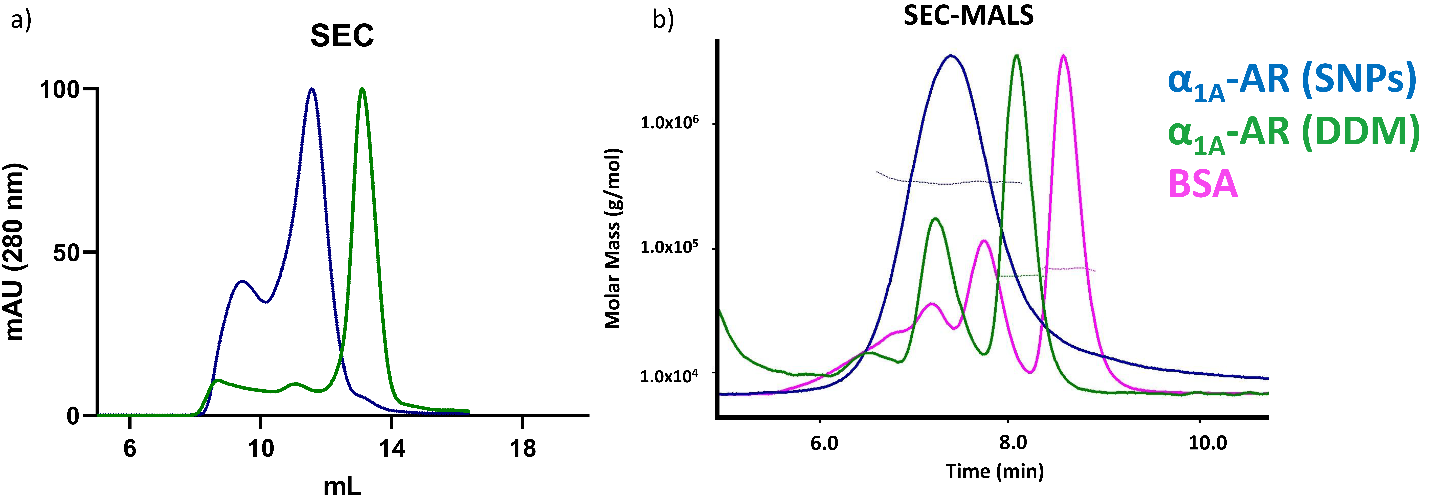

**Supplementary Figure 4. Comparing SEC and SEC-MALS profiles of α_1A_-AR in DDM and SNPs.** a) SEC profiles of α_1A_-AR in DDM (green) and small loaded SNPs (blue) where the latter were produced at 1:20:250 molar ratio of α_1A_-AR:saposin A:POPC, pH 7.4. The shorter retention time of SNPs indicates their larger hydrodynamic volume following reconstitution. The leading peak corresponds to larger loaded SNPs. b) SEC-MALS analysis of the small loaded SNPs (blue) compared with α_1A_-AR in DDM (green) and bovine serum albumin (BSA, pink). SNPs exhibited a substantially larger hydrodynamic volume than DDM incorporated receptor, while the straight line in the plots confirms acceptable homogeneity across all samples.

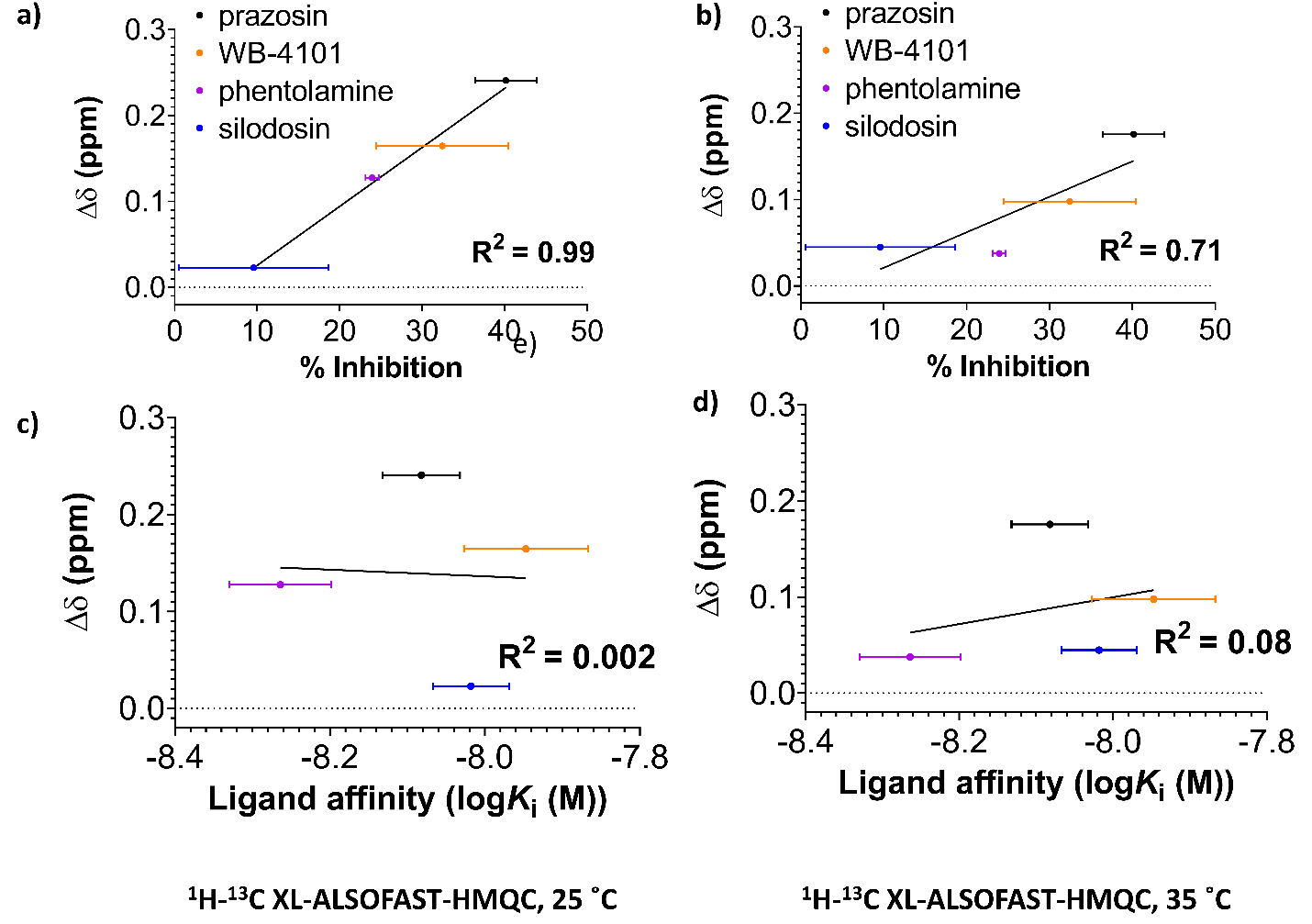

**Supplementary Figure 5. Analysis of average chemical shift differences (Δδ) of the M203^5.57^ in relation to ligand inhibitory efficacy and affinity**. The chemical shift difference (Δδ) of the ^13^C^ε^H_3_ of M203^5.57^ in α_1A_-AR A4 (L312F) were measured in the presence of prazosin (black), WB-4101(orange), phentolamine (magenta), and silodosin (blue). Δδ values were plotted against ligand inhibitory efficacies derived from previously published data on a constitutively active α_1A_-AR mutant (1). A clear linear regression was observed between the average chemical shifts of the M203^5.57^ and inhibitory efficacy in a) ^1^H,^13^C XL-ALSOFAST-HMQC spectra at 25 °C and b) 35 °C. In contrast, no correlation was found between Δδ and ligand affinities in c) 25 °C and d) 35 °C experiments. Average chemical shift differences (Δδ) were calculated as Δδ = [(Δδ_1H_)^2^ + (Δδ_13C_/3.5)^2^]^0.5^.

Supplementary Table 1. Comparing the relative signal intensities for M203^5.57^ using different 2D ^1^H-^13^C-HMQC pulse sequences and temperature.

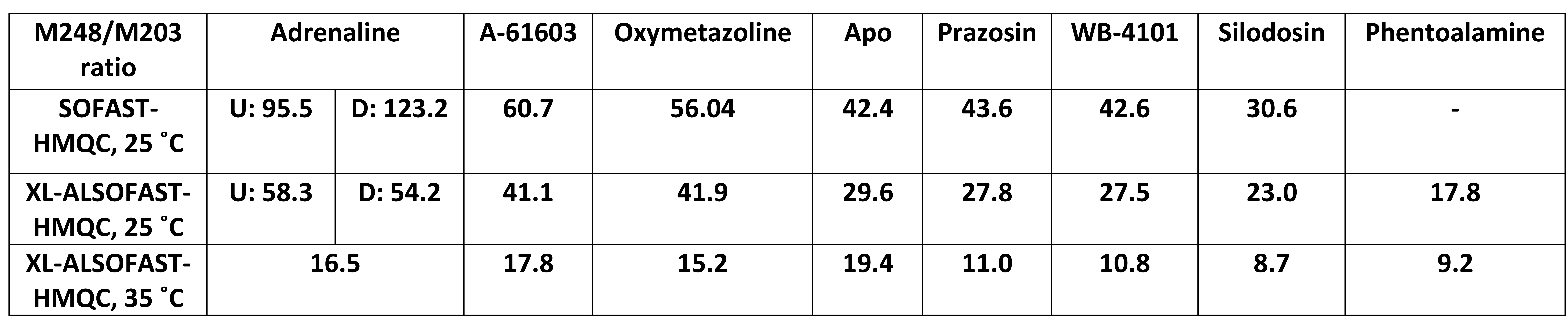

Supplementary Table2. Comparing the relative signal intensities for M115^3.41^ using different 2D ^1^H-^13^C-HMQC pulse sequences and temperature.

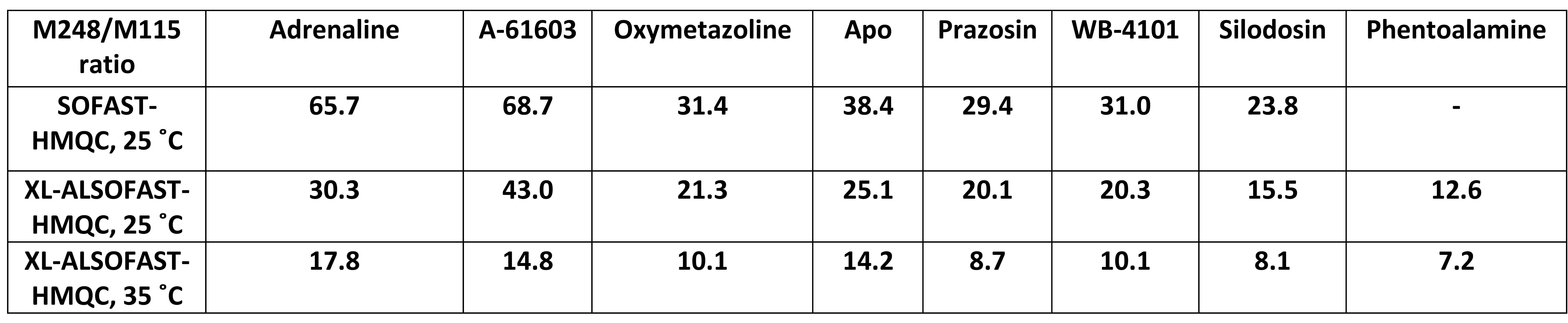

1.	Zhu J, Taniguchi T, Takauji R, Suzuki F, Tanaka T, Muramatsu I. Inverse agonism and neutral antagonism at a constitutively active alpha-1a adrenoceptor. British Journal of Pharmacology. 2000;131(3):546-52.

2.	Wu F-J, Williams LM, Abdul-Ridha A, Gunatilaka A, Vaid TM, Kocan M, et al. Probing the correlation between ligand efficacy and conformational diversity at the α1A-adrenoreceptor reveals allosteric coupling of its microswitches: Correlation of conformation with ligand efficacy for α1A-AR. Journal of Biological Chemistry. 2020;295(21):7404-17.
